# Rtapas: An R package to assess cophylogenetic signal between two evolutionary histories

**DOI:** 10.1101/2022.05.17.492291

**Authors:** Mar Llaberia-Robledillo, J. Ignacio Lucas-Lledó, Oscar Alejandro Pérez-Escobar, Boris R. Krasnov, Juan Antonio Balbuena

## Abstract

Cophylogeny represents a framework to understand how ecological and evolutionary process influence lineage diversification. However, linking patterns to mechanisms remains a major challenge. The recently developed Random Tanglegram Partitions provides a directly interpretable statistic to quantify the strength of cophylogenetic signal, maps onto a tanglegram the contribution to phylogenetic signal of individual host-symbiont associations, and can incorporate phylogenetic uncertainty into estimation of cophylogenetic signal. We introduce `Rtapas` (v1.2), an R package to perform Random Tanglegram Partitions. `Rtapas` applies a given global-fit method to random partial tanglegrams of a fixed size to identify the associations, terminals, and nodes that maximize phylogenetic congruence. `Rtapas` extends the original implementation with a new algorithm that tests phylogenetic incongruence and adds ParaFit, a method designed to test for topological congruence between two phylogenies using patristic distances, to the list of global-fit methods than can be applied. `Rtapas` can particularly cater for the need for causal inference in cophylogeny as demonstrated herein using to two real-world systems. One involves assessing topological (in)congruence between phylogenies produced with different DNA markers and identifying the particular associations that contribute most to topological incongruence, whereas the other implies analyzing the evolutionary histories of symbiont partners in a large dataset. `Rtapas` facilitates and speeds up cophylogenetic analysis, as it can handle large phylogenies reducing computational time, and is directly applicable to any scenario that may show phylogenetic congruence (or incongruence).

---

One of the main challenges in evolutionary biology is inferring how extant interactions within ecological communities translate into species diversification. (Cavender-Bares et al. 2009; Poisot 2015; Blasco-Costa et al. 2021). Cophylogeny provides a framework to understand how ecological and evolutionary process influence lineage diversification (Hutchinson et al. 2017; Blasco-Costa et al. 2021). When organisms are linked through interspecific interactions over long evolutionary time (such as host-parasite or plant-pollinator interactions), their diversification is rarely independent (de Vienne et al. 2013; Allio et al. 2021; Fuzessy et al. 2022). As the phylogenies of associated taxa are often dependent, a certain degree of topological similarity between them is expected (Balbuena et al. 2013). Thus, phylogenetic congruence quantifies this similarity by the extent to which each node and branch-length in a phylogenetic tree maps to a corresponding position in the other phylogenetic tree (de Vienne et al. 2007). Perfect congruence could be an indicator of cospeciation while a total absence of congruence would indicate random associations in the evolutionary history of the taxa involved (Legendre et al. 2002; Balbuena et al. 2013). However, there are other types of evolutionary events that can produce some degree of topological congruence between two phylogenies, such as preferential host-switching events (Charleston and Robertson 2002) or pseudocospeciation (de Vienne et al. 2013). Thus, linking patterns with mechanisms probably is the main outstanding challenge in cophylogeny (Blasco-Costa et al. 2021).

Recently, Balbuena et al. (2020) introduced Random Tanglegram Partitions (Random TaPas), a novel approach that provides a directly interpretable statistic to quantify the strength of cophylogenetic signal and maps the contribution to phylogenetic signal of individual host-symbiont associations onto a tanglegram. Random TaPas is based on the repeated random subsampling of the set of associations (or interactions) between taxa from the two phylogenies. It produces a Gini coefficient, either conventional (*G*) or normalized (*G*\*) (Raffinetti et al. 2015), that measures how evenly distributed the cophylogenetic signal is along the tanglegram. It has been shown that *G\** is negatively correlated with the amount of cospeciation events in simulated data sets (Balbuena et al. 2020). Thus, low values of *G\** suggest a strong cophylogenetic signal across the whole dataset, while high values of *G\** may indicate either cophylogenetic signal concentrated in parts of the tanglegram or low overall cophylogenetic signal. Random Tapas is especially helpful in aiding the biological interpretation of large tanglegrams (e.g. Dupeyron et al. 2021; Léveillé-Bourret et al. 2021) and provides a way to incorporate phylogenetic uncertainty into estimation of cophylogenetic signal (e.g. Hayward et al. 2021).

Here we develop Random TaPas as an R package, `Rtapas` (v1.2) that extends the original implementation by adding a new algorithm that scans the host-symbiont tanglegram for phylogenetic incongruence. It also incorporates ParaFit (Legendre et al. 2002) as global-fit method to implement Random TaPas, in addition to geodesic distances (GD) (Schardl et al. 2008) and PACo (Balbuena et al. 2013) originally implemented in Balbuena et al. (2020). `Rtapas` further enhances the usability and implementation of Random TaPas by including functions (i) to facilitate the prior processing of association data between taxa, (ii) to estimate, in a set of probability trees, the statistic of a given global-fit method, (iii) to estimate the (in)congruence metrics of the individual host-symbiont associations, and (iv) to compute either *G* or *G\** characterizing the distribution of such metrics (Table 1). In addition, the procedures to tackle phylogenetic uncertainty have been enhanced by incorporating new arguments to optionally parallelize the processes. This allows to relieve the excessive computational load with large datasets.

**Table 1.** Overview of the Rtapas functions

| Function | Description |
| --- | --- |
| `assoc_mat()` | Creates a binary host-symbiont association matrix from a two-column matrix representing associations between host and symbiont species. |
| `one2one_f()` | For a binary matrix of host-symbiont associations, it finds the maximum number of host-symbiont pairs, $n$, for which one-to-one unique associations can be chosen. |
| `trimHS_maxC()` | Chooses randomly $n$ unique one-to-one associations for $N$ runs and trims the host-symbiont association matrix to include only the $n$ associations. |
| `trimHS_maxI()` | Chooses randomly $n$ associations for $N$ runs and trims the host-symbiont association matrix to include only the $n$ associations. |
| `max_cong()` | Prunes the host ($H$) and symbiont ($S$) phylogenies to conform with the trimmed matrix and computes the given global fit method, Geodesic Distances (GD), Procrustes Approach to Cophylogeny (PACo) or ParaFit (Legendre et al. 2002) between the pruned trees. Then, determines the frequency of each host-symbiont association occurring in a given percentile of cases that maximize phylogenetic congruence. |
| `max_incong()` | Prunes $H$ and $S$ to conform with the trimmed matrix and computes the given global fit method (PACo or ParaFit) between the pruned trees. Then, determines the frequency of each host-symbiont association occurring in a given percentile of cases that maximize phylogenetic incongruence. |
| `prob_statistic()` | Computes frequencies of the host-symbiont associations for pairs ($H$ and $S$) of posterior probability trees from the statistics generated with GD, PACo or ParaFit. |
| `linkf_CI` | Computes confidence intervals of the estimated (in)congruence metrics of the individual host-symbiont associations using a set of pairs ($H$ and $S$) of posterior probability trees. |
| `tangle_gram()` | Maps the estimated (in)congruence metrics of the individual host-symbiont associations as heatmap on a tanglegram. |
| `gini_plot()` | Computes and plots Gini coefficients and their confidence intervals of the frequency distributions of the estimated (in)congruence metrics of the individual host-symbiont associations. |

Herein we provide an overview of the new R package and further demonstrate its capabilities with two real-world examples each representing a different major domain of application of cophylogenetic analysis.

## Overview of rtapas

`Rtapas` requires a triple as data input consisting of the host and symbiont phylogenies (*H* and *S* respectively) and a binary matrix (**A**) encapsulating the associations between *H* and *S* (Balbuena et al. 2020). The triple is often represented as a tanglegram, in which *H* and *S* are displayed face to face, and the associations in **A** are represented as lines connecting terminals in *H* and *S*. The input parameters of the original Random TaPas algorithm are the number of one-to-one unique associations to be sampled randomly (*n*), the number of replicates to compute a global-fit statistic with subtanglegrams of *n* associations (*N*) and a percentile (*p*) of the distribution of the *N* statistics generated. `Rtapas` also includes a new algorithm that identifies host-symbiont associations that maximize incongruence between the two phylogenies (Fig. 1). The main difference with respect to the original Random TaPas algorithm is that the *n* host-symbiont associations chosen in each run are no longer required to be one to one, that is, multiple associations in any subtanglegram are allowed (Fig. 1).

**Figure 1.**
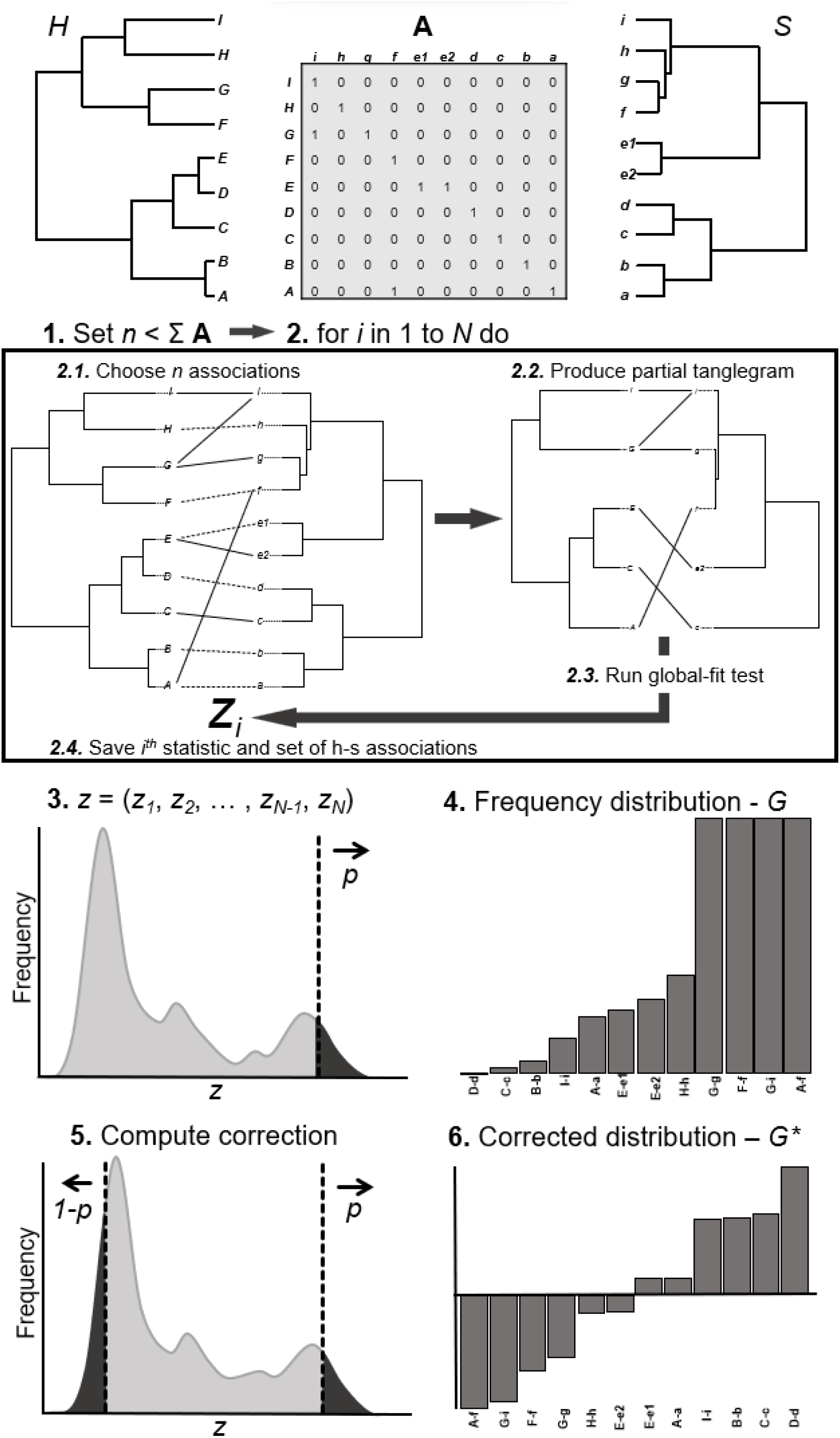
Stepwise methodology for the new maximize incongruence algorithm. Given a triple with the phylogenies of hosts (*H*) and symbionts (*S*), and a binary matrix (**A**) that codes the associations between terminals in *H* and *S*: (1) Set a number *n* less than the total number of host–symbiont associations. (2) For *i* from 1 to *N* times (where *N* is typically ≥ 10^4^) do (2.1) Randomly choose *n* associations in **A** allowing multiple associations; (2.2) Produce a partial tanglegram that includes only the *n* associations chosen at Step 2.1 by trimming **A** and pruning *H* and *S*; (2.3) Run a global-fit test with the partial triple and (2.4) Save the resulting statistic *z*_*i*_ and the set of *n* host–symbiont associations selected at 2.1. (3) Render the frequency distribution of the *z*_*i*_’s and set a percentile *p* where the highest cophylogenetic incongruence is expected. (In this example, *z*_*i*_ is expected to be directly proportional to incongruence). (4) Determine how many times each host–symbiont association occurs in *p*, return the distribution of observed frequencies, and compute a Gini coefficient (*G*). (5) Optionally in systems in which a large number of host-parasite associations are similar in either their contribution to (in)congruence, the incongruence of each association can be computed as the difference of frequencies of occurrence in the *p* and *1-p* percentiles. (6) The distribution of incongruence values is returned and a normalized Gini coefficient (*G∗*) measuring its shape is produced.

### Original Algorithm

We illustrate the application of `Rtapas` applying the original algorithm to a dataset of the trematode *Coitocaecum parvum* (Opecoelidae) and its amphipod host *Paracalliope fluviatilis* (Paracalliopiidae) from seven locations in South Island, New Zealand (Lagrue et al. 2016; Balbuena et al. 2022). We first load the library and the dataset:

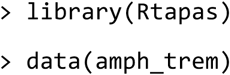

Typically, the number of replicates to be run (*N*) is set to 10,000, as higher values can be computationally prohibitive.

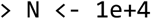

In certain configurations, setting the number of unique host-associations (*n*) requires some work because there is an upper limit to the number of such associations that can be chosen in any of the *N* runs. Function `one2one_f` determines the maximum *n* for which one-to-one associations can be chosen in all *N* runs. The range of *n*s tested (from 5 to 15 in the example below) is specified with `interval`.

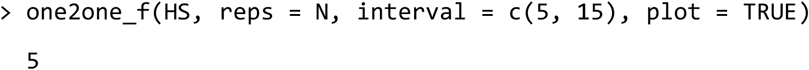

The output indicates that 5 (or 6 in some cases depending on the randomization) is such maximum *n* value. Balbuena et al. (2020) recommend using an *n* between 10 and 20 % of the total number of associations in **A**, i.e. 75 in the present example. Thus *n* = 5 is not ideal.

If the `plot` argument is set to `TRUE` (the default), a plot of *n* in the `interval` range against the number of runs that could be completed is produced (Fig. 2). Based on this information, setting *n* = 8 offers a good compromise between the number of unique runs implemented (> 9,500) and the fraction (> 10%) of total number of host-symbiont associations recommended for analysis. Thus,

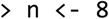

**Figure 2.**
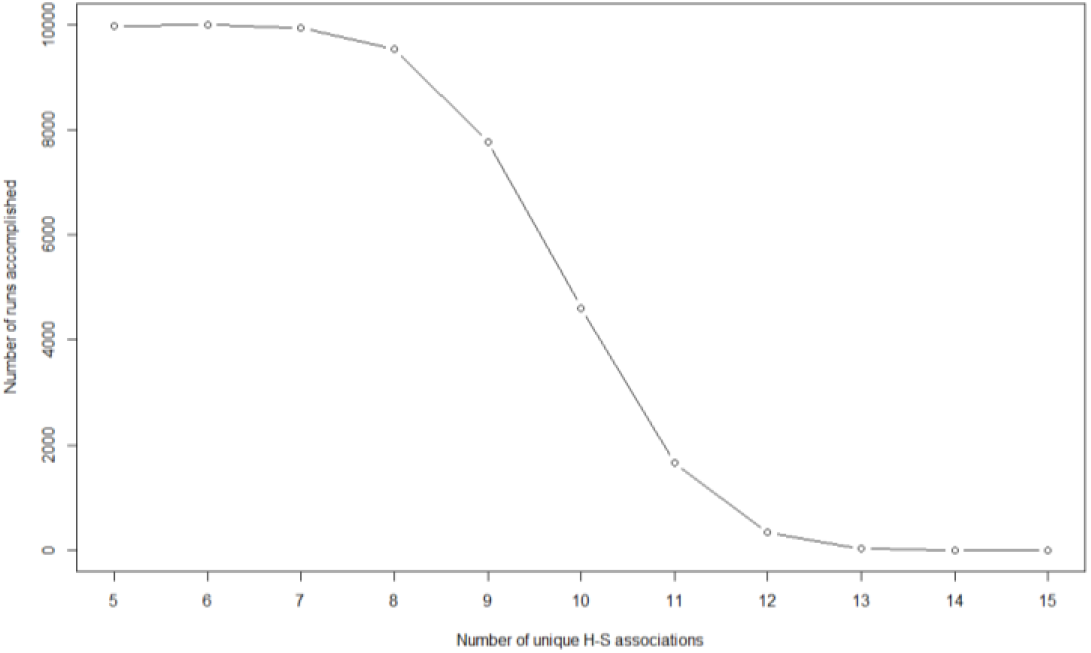
Number of runs (*N*) of the original Random TaPas algorithm that can be accomplished with a varying number of host-symbiont associations (*n*) in the trematode *Coitocaecum parvum* with its amphipod host, *Paracalliope fluviatilis* example.

Next, `max_cong` is applied to identify host-symbiont associations that maximize congruence between two phylogenies, in three main steps. First, for *N* runs, *n* unique associations in **A** are selected at random. Then, a list of the *N* trimmed matrices is produced and a global-fit method (the user can choose among PACo, as in the example below, GD and ParaFit) is applied to each trimmed matrix, i.e. only considering the *n* host-symbiont associations selected. This yields a vector of *N* statistics produced by the global-fit method of choice. Since the three methods available produce statistics inversely proportional to cophylogenetic congruence, the most congruent host-symbiont associations would correspond to statistics on the left tail of their distribution. The last step is to determine the frequency of occurrence of each host-symbiont association in a *p* (`percentile = 0.01` by default) of the distribution of *N* statistics generated.

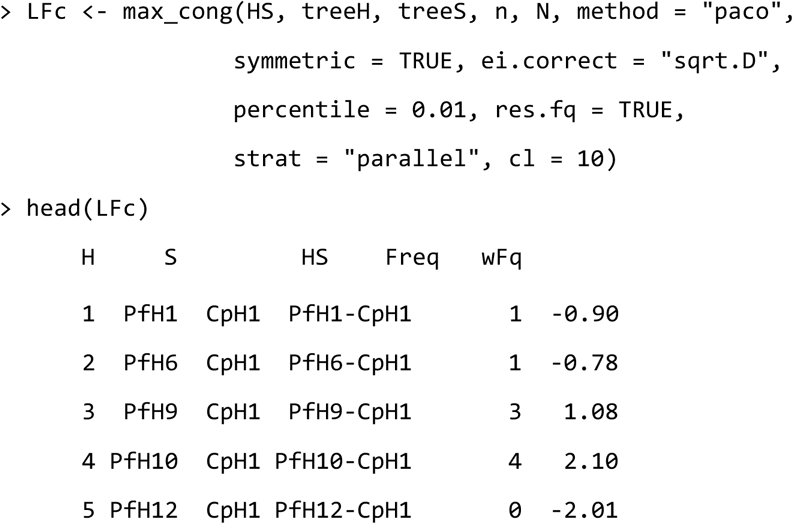

The output `LFc` is a data frame with the names of the terminals, their combinations based on the interactions and the frequency of occurrence of the association (`Freq`) (the R base function `head` is used above to show the first lines of the output). In addition, a corrected residual frequency (`wFq`) is also computed if `res.fq = TRUE`. This correction is required in tanglegrams with multiple associations (i.e. one host associated with two or more symbionts or vice versa) because the algorithm biases for one-to-one associations in the overall frequency distribution (Balbuena et al. 2020).

### New Algorithm

Function `max_incong` implements the new algorithm to identify host-symbiont associations that are most disruptive of congruence between *H* and *S*. We illustrate its application to the same trematode-amphipod dataset. The steps are analogous to those in `max_cong`. However, since the global-fit methods contemplated produce statistics that are directly proportional to incongruence, the frequency of occurrence of each host-symbiont association are evaluated in a high *p* (`percentile = 0.99` is the default) (Fig. 1). This algorithm is less restrictive in terms of the *n* chosen as it allows multiple associations and it internally controls for every subtree (Fig. 1) to have at least three terminals. Below we also use *n* = 8 as in the previous `max_cong` analysis:

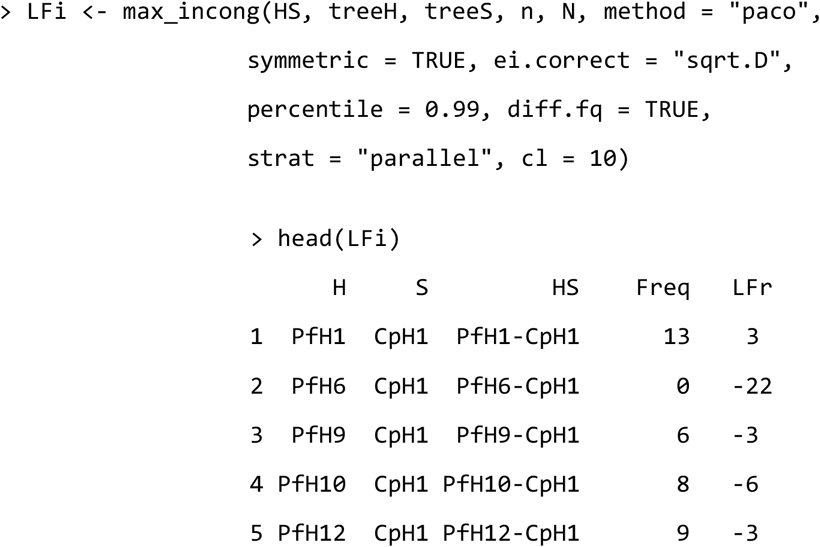

In the output `LFi`, the incongruence of each association is measured as the frequency of occurrence of a given association in the *p* of the distribution (`Freq`). However, in systems in which a large number of host-parasite associations are similar in either their contribution to congruence or incongruence, a given association can occur with certain frequency at both the 0.01 and 0.99 percentiles. We recommend in these scenarios setting `diff.fq = TRUE`. So, the incongruence of the association (`LFr` above) is estimated as *f*_*1-p*_ – *f*_*p*_, where *f*_*p*_ and *f*_*1-p*_ are the frequency of occurrence of the association in the *p* and 1 – *p* percentiles respectively (Fig. 1). This adjustment makes comparison with residual frequencies produced by the original algorithm (`wFq`) straightforward, as `wFq` and `Lfr` are directly proportional to each other.

### Graphical Output

We use `tangle_gram` to display the quantitative information in `LFc` and `LFi` on a tanglegram (Fig. 3a, b). The raw (`Freq`) and the adjusted (either `wFr` or `Lfr`) frequencies represent estimates of topological congruence (or incongruence) of each host-symbiont association. These estimates can be interpreted as contributions to the global cophylogenetic signal that are displayed as a heatmap on the tanglegram. The argument `colgrad` defines the color scale.

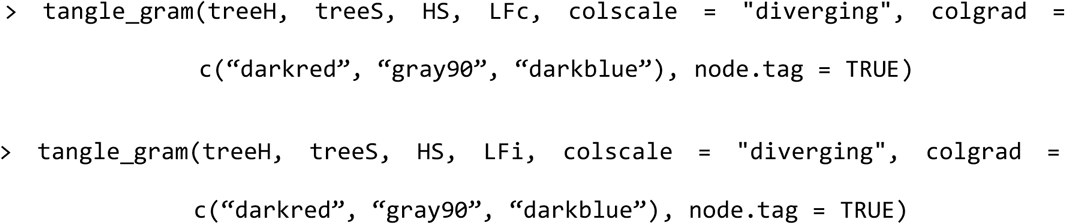

**Figure 3.**
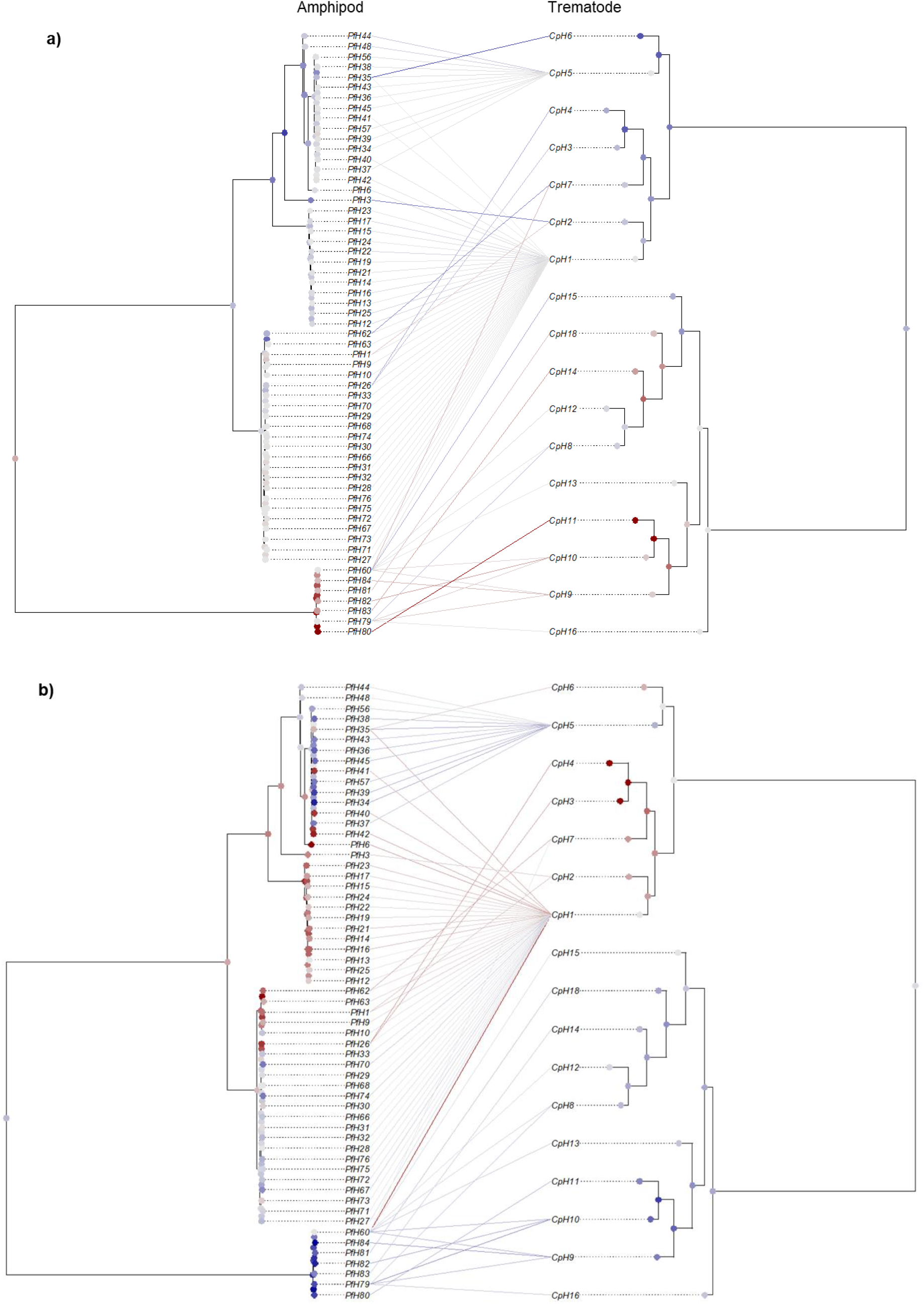
Tanglegrams representing the associations between the trematode *Coitocaecum parvum* with its amphipod host, *Paracalliope fluviatilis*. (a) Residual frequency corresponding to each trematode-amphipod association obtained with the algorithm that identifies associations that maximize congruence. (b) Corrected frequency corresponding to each trematode-amphipod association obtained with the algorithm that identifies associations that maximize incongruence. Both algorithms were applied with PACo. The frequencies are mapped using a color scale centered at light grey (zero) ranging from dark red (lowest/incongruent) to dark blue (highest/congruent). The average residual frequency of occurrence of each terminal and fast maximum likelihood estimators of ancestral states of each node are also mapped according to the same scale.

If `res.fq = TRUE` (`max_cong`) or `diff.fq = TRUE` (`max_incong`), the metrics to be plotted include positive and negative values. In such a case, we recommend to set the argument `colscale = “diverging”`, as it centers the color scale at 0 and displays negative and positive values along two respective color gradients. Otherwise, with `colscale = “sequential”`, the scale spans a single-color gradient. Furthermore, if `node.tag = TRUE`, the average `LFc` and `LFi` output value of each terminal is likened to a continuous trait, and estimators of ancestral states can be computed and displayed at the nodes of the phylogeny (Fig. 3). This procedure is useful to assess different levels of cophylogenetic signal across *H* and *S*. The estimation is performed with `fastAnc` of package `phytools` (Revell 2012).

### Assessing Phylogenetic Uncertainty

An improved aspect in `Rtapas` is the simplification in assessing the influence of phylogenetic uncertainty on the estimates obtained with `max_cong` and `max_incong`. First, `trimHS_maxC` (`max_cong` algorithm) or `trimHS_maxI` (`max_incong` algorithm) are used to produce a list of *N* trimmed matrices. For instance,

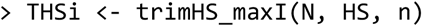

Then, a global-fit statistic is computed for a given set of pairs (*H* and *S*) of the posterior probability trees (`freqfun = “paco”` below indicates that we chose PACo’s *m*^2^):

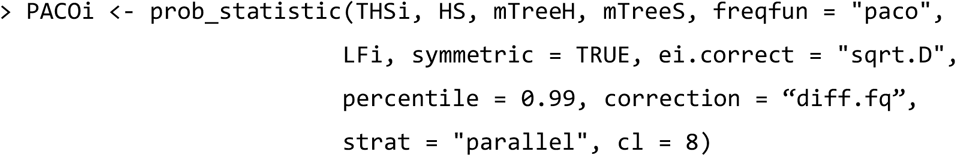

where `mTreeH` and `mTreeS` are 1,000 posterior probability trees of hosts and symbionts, respectively. Next, the contribution of each host-symbiont association to (in)congruence are estimated and displayed as a bar graph (Supplementary Material, Fig. S1):

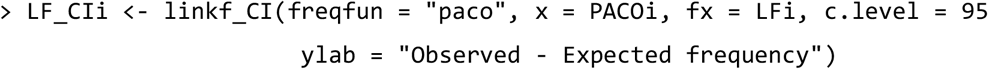

### Gini Coefficients

Finally, the shape of the distribution of the contributions of the host-symbiont associations to (in)congruence can be characterized with a Gini coefficient, either *G* or *G*\* (Raffinetti et al. 2015) if negative values are produced due to corrected frequencies (either if `res.fq = TRUE` or `diff.fq = TRUE`). Since both residuals and corrected frequencies have been estimated here (applying PACo), the *G*\* values of the consensus trees, with its confidence intervals obtained with the set of pairs of posterior probability trees, are estimated (Fig. 4):

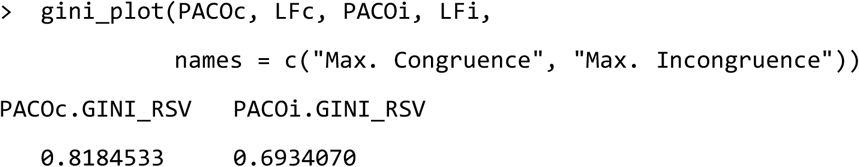

**Figure 4.**
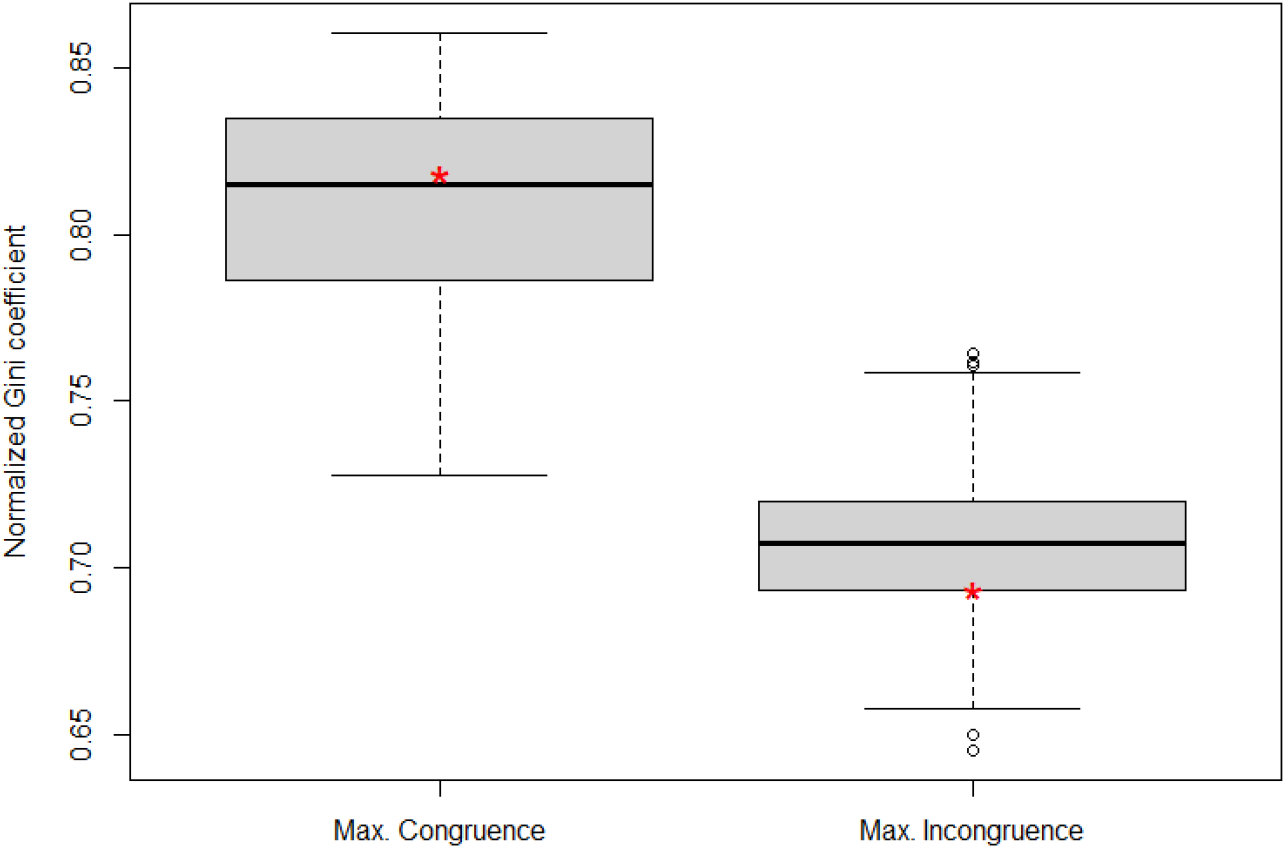
Normalized Gini coefficient (*G\**) of the frequency distributions applying `max_cong` and `max_incong` algorithms with PACo to the consensus trees (asterisk) and to 1,000 randomly chosen pairs posterior probability trees (boxplots).

The Gini value is inversely proportional to cophylogenetic signal. If the distribution of congruence (or incongruence) across the host-symbiont associations were random, the expected value of *G*\* would be 2/3 (or 1/3 if *G* is employed). Values lower than these thresholds would indicate higher cophylogenetic signal than expected by chance (Balbuena et al. 2020).

## Case studies

To demonstrate the usability of `Rtapas`, we present two case studies in two domains of applicability.

### Phylogenetic congruence of chloroplast and nuclear loci of orchids

For decades, plastid loci have been used for phylogenetic inference due to the high abundance of organellar DNA in a cell (Bateman et al. 2021). However, the plastid and nuclear genomes often differ in the rates of substitution of nuclear loci (Smith et al. 2014), their modes of inheritance (Gonçalves et al. 2020) and replication (Heinhorst and Cannon 1993), which can lead to divergent and incongruent evolution between topologies (Pérez-Escobar et al. 2016, 2017, 2021; Rose et al. 2021). Here, we examined the congruence between the nuclear and chloroplast phylogenies (consensus and 1,000 pairs of posterior probability trees) originally analysed by Pérez-Escobar et al. (2021), consisting of 52 orchid genera. We designated the nuclear data as *H*, and the plastid data as *S*. We chose PACo as the global-fit method, with `symmetric = FALSE` (the default), i.e. we assumed that that the evolutionary history of the chloroplasts tracks to some extent that of the nuclear genome (Pérez-Escobar et al. 2016). We displayed the phylogenies in two tanglegrams (Supplementary Material, Fig. S2) and represented the frequency of occurrence of the plastid–nuclear associations (`Freq` in `LFc` and `LFi`) and their 95% confidence intervals derived from the comparison 1,000 pairs of posterior probability trees (Fig 5a, b).

**Figure 5.**
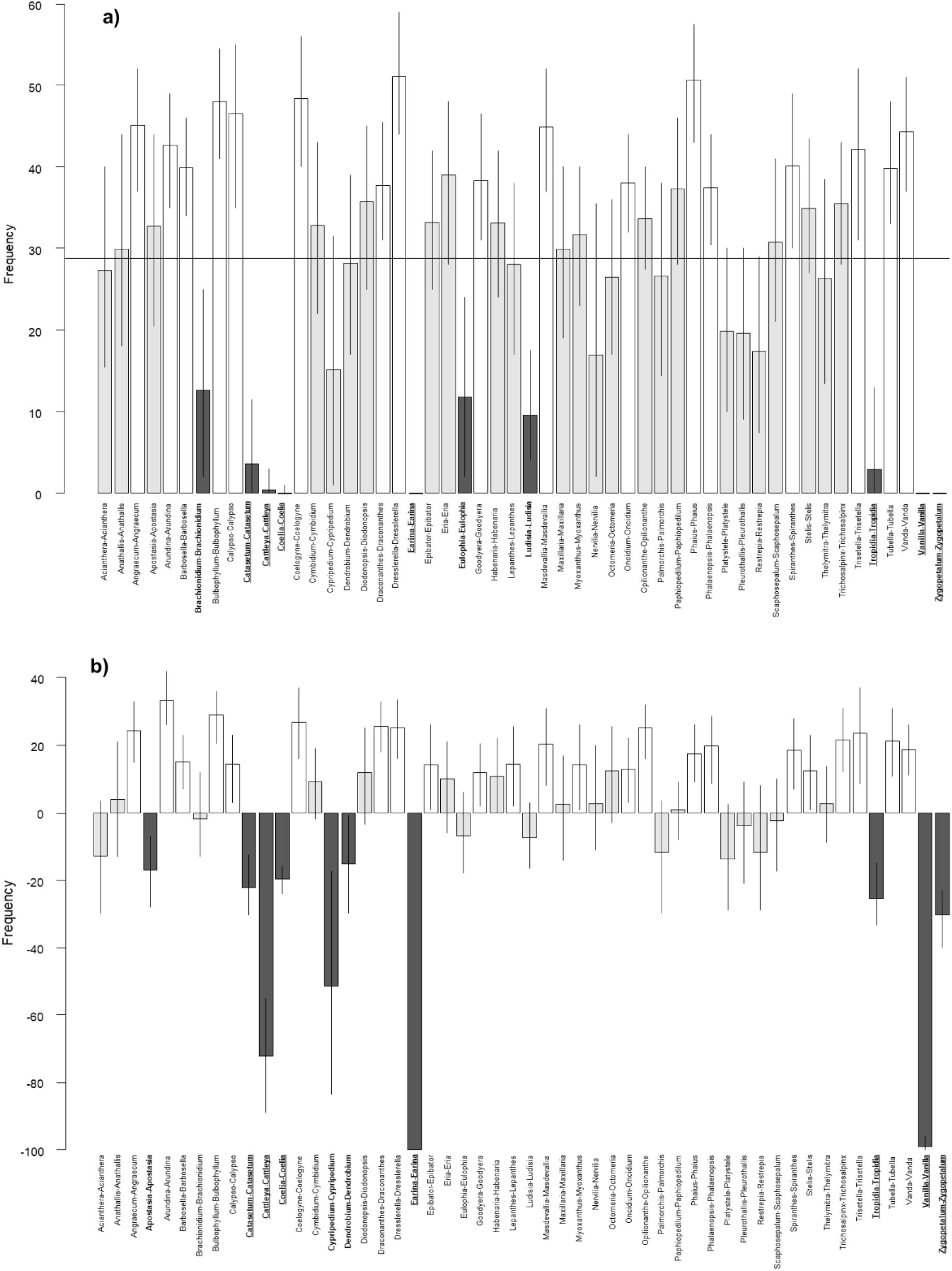
(a) Average of frequency distributions for maximizing congruence algorithm. (b) Corrected frequency distributions for maximizing incongruence algorithm chloroplast and nuclear loci. Both algorithms were applied with PACo. Vertical lines represent 95% confidence intervals of the frequencies computed empirically with 1,000 randomly chosen pairs of posterior probability trees. The white bars represent those associations with higher congruence, the gray bars the neutral associations, and the black bars the associations with higher incongruence. Terminals in bold and underlined are those identified as incongruent by both algorithms (`max_cong` and `max_incong`).

At least 10 terminals showed conflicting positions (incongruence). Both algorithms agreed at identifying seven of the topological mismatches, except for three terminals that only appear as conflicting in one of the both algorithms (`max_cong` and `max_incong`). Eight of these terminals, *Cattleya, Coelia, Earina, Catasetum, Eulophia, Tropidia* and *Vanda*, matched the conflicting positions identified by Pérez-Escobar et al. (2021). Interestingly, two of these terminals, *Vanda* and *Zygopetalum*, belong to two subtribes, Aeridinae and Zygopetalinae, respectively, whose phylogenetic positions are incongruent at deeper phylogenetic levels (Pérez-Escobar et al. 2021). Additionally, deeper branches linking *Zygopetalum* and *Vanda* to their MRCA (i.e., Cymbideae + Vandeae) showed labile positions in the sets of nuclear and plastid posterior probability trees (see Appendix S10 in Pérez-Escobar et al. 2021) which could account for being flagged as incongruent by `Rtapas`. Thus, the incorporation of posterior probability trees may reveal patterns that may go unnoticed considering only a pair of consensus trees.

### Phylogenetic congruence between small mammals and their flea parasites

`Rtapas` can also unravel complex patterns in large host-symbiont system. We use a dataset from Krasnov et al. (2016) consisting of 130 species of small mammals and 202 species of flea parasites. Both `max_cong` and `max_incong` were applied with multiple association correction (`res.fq = TRUE`) and subtraction of frequencies in extreme percentiles (`diff.fq = TRUE`), respectively. We chose PACo as the global-fit method with `symmetric = FALSE`, as we assumed that flea diversification is driven by that of the host.

Both algorithms rendered similar results. In mammals, Eulipothyphla (Talpidae and Soricidae) and ochotonid Lagomorpha displayed higher incongruence (nodes and terminals in red) (Fig. 6, Supplementary Material Fig. S3). By contrast, Rodentia displayed higher congruence (marked in blue), except the suborder Sciuromorpha whose associations with fleas were mostly phylogenetically incongruent (Fig. 6). Fleas exhibited a far more irregular pattern, with terminals and nodes varying greatly in congruence and incongruence within clades. Thus, the degree of cophylogenetic incongruence in hosts was concentrated in clades, whereas it was distributed across the entire tree in fleas. This suggests that within a flea family some clades exhibit some phylogenetic conservatism in relation to their hosts, whereas other clades do not, due probably to host switching.

**Figure 6.**
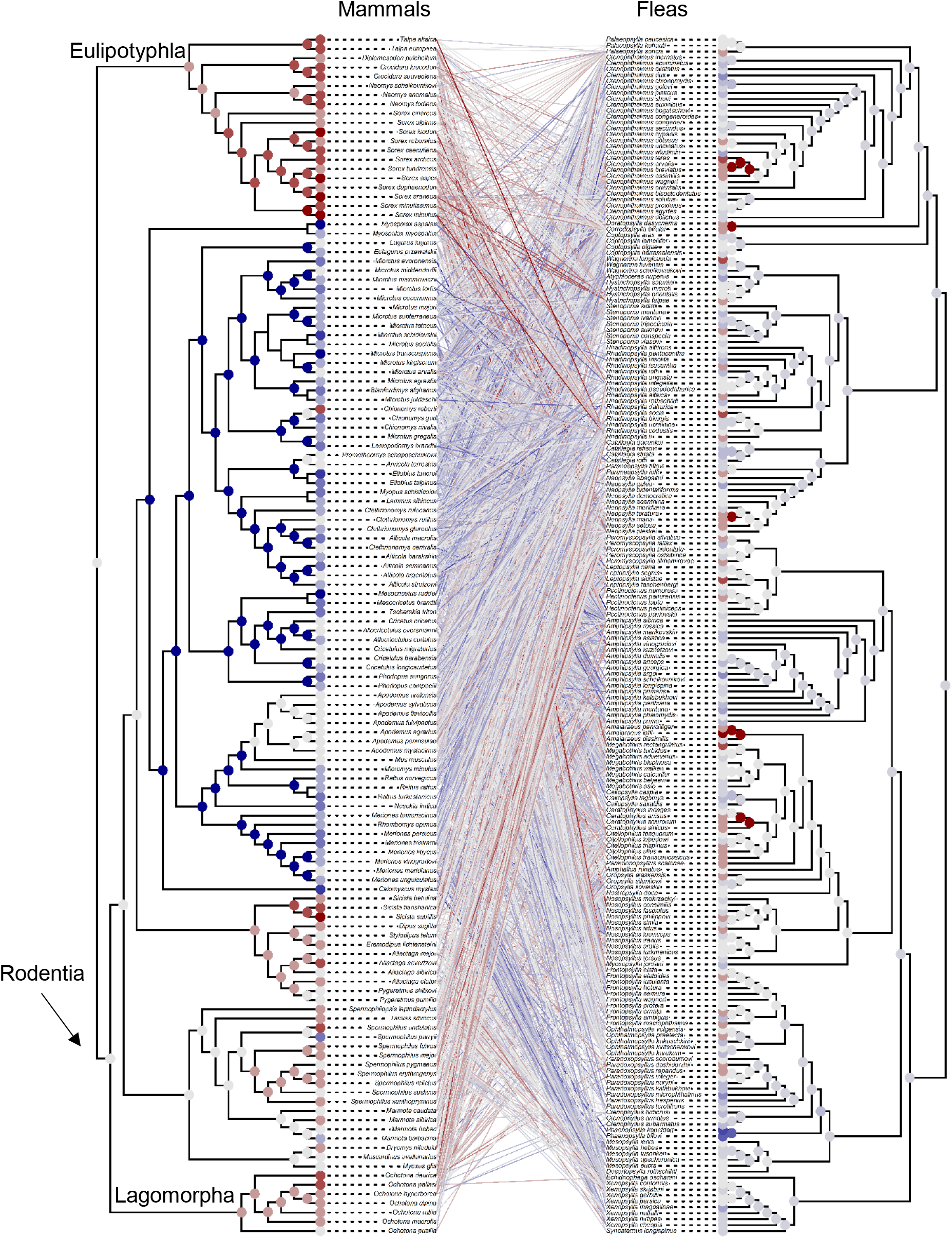
Tanglegram representing the associations between the mammal species with their flea parasite species with the algorithm to maximize incongruence using PACo. The corrected frequenciescorresponding to each mammal-flea association obtained are mapped using a color scale centered at light grey (zero) ranging from dark red (lowest/incongruent) to dark blue (highest/congruent). The average residual frequency of occurrence of each terminal and fast maximum likelihood estimators of ancestral states of each node are also mapped according to the same scale.

These results not only support Krasnov et al.’s (2016) findings, but also provide new information by flagging those taxa that exhibit a greater cophylogenetic signal. Since host– symbiont associations are marked as congruent or incongruent relative to the other associations in the triple (Balbuena et al. 2020), prospective users can also run partial analyses within designated clades, as illustrated in Supplementary Material (Fig. S4, S5a, S5b, S6), to gain more detailed insight and lower taxonomic levels.

## Final remarks

As exemplified by the case studies presented, we posit that `Rtapas` can improve causal inference in cophylogeny in two domains. First, `Rtapas` provides a straightforward procedure to assess topological (in)congruence between phylogenies produced with different DNA markers and specifically identify the associations that contribute most to topological incongruence. This issue is of paramount importance in evolutionary biology, since identification of these outlier terminals can shed light onto evolutionary events and processes such as horizontal gene transfer (Bellot and Renner 2016), gene flow (Vargas et al. 2017), and incomplete lineage sorting (de Vienne et al. 2012; Knowles et al. 2018). An alternative to analyzing entire genomes (e.g., Pérez-Escobar et al. 2021) is to identify a subset of loci for phylogenetic analysis that are most likely to reflect gene-tree evolutionary histories that are congruent with the most frequent/dominant species tree relationships (e.g., Bogarín et al. 2018; Renner et al. 2021). So `Rtapas` can critically contribute to identify coalescent data partitions.

Second, the expansion of DNA sequencing and phylogenetic reconstruction has resulted in increasingly common scenarios involving large trees (e.g., 100+ terminals; Eiserhardt et al. 2018). `Rtapas` can handle such complex datasets and deliver insightful information on the shared evolutionary histories. For instance, our analysis provides strong evidence for an asymmetrical diversification of mammals and their flea parasites. In addition, metrics of cophylogenetic signal rendered by `Rtapas` allow assessing to which extent extant associations among species have been the product of a coupled evolutionary history.

Beyond testing phylogenetic congruence, `Rtapas` is amenable to any problem requiring the comparison of dissimilarity matrices. Thus, it could be a most appropriate method to implement the Cophylospace approach recently proposed by Blasco-Costa et al. (2021) to enhance the explanatory power of cophylogenetic analysis. This framework relies on the cross assessment of morphological and phylogenetic similarities between hosts and symbionts. `Rtapas` could play a crucial role at identifying differences among clades on the intensity of coevolutionary and phylogenetic tracking between symbiotic partners.

## Acknowledgements

This work was supported by MCIN/AEI/10.13039/501100011033 (PID2019-104908GB-I00), Government of Spain. MLR is supported by a pre-doctoral contract from MCIN/AEI/10.13039/501100011033 (Government of Spain) and by “European Union Next Generation EU/PRTR (PRE2020-095070). OAPE acknowledges support from the Sainsbury Orchid Fellowship at the Royal Botanic gardens, Kew, and the Swiss Orchid Foundation.

## Data accessibility

The development version of `Rtapas` is on Github at https://github.com/mllaberia/Rtapas. The code necessary to reproduce the analyses is on Github at https://github.com/mllaberia/Rtapas/…, and an archived version is accessible on Zenodo https://doi.org/X

## Notes

https://github.com/mllaberia/Rtapas

### Competing Interest Statement

The authors have declared no competing interest.

## References

Allio R., Nabholz B., Wanke S., Chomicki G., Pérez-Escobar O.A., Cotton A.M., Clamens A.-L., Kergoat G.J., Sperling F.A.H., Condamine F.L. 2021. Genome-wide macroevolutionary signatures of key innovations in butterflies colonizing new host plants. Nat Commun. 12:354.

Balbuena J.A., Míguez-Lozano R., Blasco-Costa I. 2013. PACo: A Novel Procrustes Application to Cophylogenetic Analysis. PLoS One. 8:e61048.

Balbuena J.A., Pérez-Escobar Ó.A., Llopis-Belenguer C., Blasco-Costa I. 2020. Random Tanglegram Partitions (Random TaPas): An Alexandrian Approach to the Cophylogenetic Gordian Knot. Syst. Biol. 69:1212–1230.

[dataset]* Balbuena J.A., Pérez-Escobar Ó.A., Llopis-Belenguer C., Blasco-Costa I. 2022. User’s Guide Random Tanglegram Partitions V.1.0.0. Zenodo. DOI: 10.5281/zenodo.6327235

Bateman R.M., Rudall P.J., Murphy A.R.M., Cowan R.S., Devey D.S., Peréz-Escobar O.A. 2021. Whole plastomes are not enough: phylogenomic and morphometric exploration at multiple demographic levels of the bee orchid clade Ophrys sect. Sphegodes. Journal of Experimental Botany. 72:654–681.

Bellot S., Renner S.S. 2016. The Plastomes of Two Species in the Endoparasite Genus Pilostyles (Apodanthaceae) Each Retain Just Five or Six Possibly Functional Genes. Genome Biol. Evol. 8:189–201.

Blasco-Costa I., Hayward A., Poulin R., Balbuena J.A. 2021. Next-generation cophylogeny: unravelling eco-evolutionary processes. Trends Ecol. Evol. 36:907–918.

Bogarín D., Pérez-Escobar O.A., Groenenberg D., Holland S.D., Karremans A.P., Lemmon E.M., Lemmon A.R., Pupulin F., Smets E., Gravendeel B. 2018. Anchored hybrid enrichment generated nuclear, plastid and mitochondrial markers resolve the Lepanthes horrida (Orchidaceae: Pleurothallidinae) species complex. Mol. Phylogenet. Evol. 129:27–47.

Cavender-Bares J., Kozak K.H., Fine P.V.A., Kembel S.W. 2009. The merging of community ecology and phylogenetic biology. Ecology Letters. 12:693–715.

Charleston M.A., Robertson D.L. 2002. Preferential Host Switching by Primate Lentiviruses Can Account for Phylogenetic Similarity with the Primate Phylogeny. Syst. Bio. 51:528–535.

de Vienne D.M., Giraud T., Shykoff J.A. 2007. When can host shifts produce congruent host and parasite phylogenies? A simulation approach. J. Evol. Biol. 20:1428–1438.

de Vienne D.M., Ollier S., Aguileta G. 2012. Phylo-MCOA: A Fast and Efficient Method to Detect Outlier Genes and Species in Phylogenomics Using Multiple Co-inertia Analysis. Mol. Biol. Evol. 29:1587–1598.

de Vienne D.M., Refrégier G., López-Villavicencio M., Tellier A., Hood M.E., Giraud T. 2013. Cospeciation vs host-shift speciation: methods for testing, evidence from natural associations and relation to coevolution. New Phytologist. 198:347–385.

Dupeyron M., Baril T., Hayward A. 2021. Broadscale evolutionary analysis of eukaryotic DDE transposons. :2021.09.26.461848.

Eiserhardt W.L., Antonelli A., Bennett D.J., Botigué L.R., Burleigh J.G., Dodsworth S., Enquist B.J., Forest F., Kim J.T., Kozlov A.M., Leitch I.J., Maitner B.S., Mirarab S., Piel W.H., Pérez-Escobar O.A., Pokorny L., Rahbek C., Sandel B., Smith S.A., Stamatakis A., Vos R.A., Warnow T., Baker W.J. 2018. A roadmap for global synthesis of the plant tree of life. Am. J. Bot. 105:614–622.

Fuzessy L., Silveira F.A.O., Culot L., Jordano P., Verdú M. 2022. Phylogenetic congruence between Neotropical primates and plants is driven by frugivory. Ecol. Lett. 25:320–329.

Gonçalves D.J.P., Jansen R.K., Ruhlman T.A., Mandel J.R. 2020. Under the rug: Abandoning persistent misconceptions that obfuscate organelle evolution. Mol. Phylogenet. Evol. 151:106903.

Hayward A., Poulin R., Nakagawa S. 2021. A broadscale analysis of host-symbiont cophylogeny reveals the drivers of phylogenetic congruence. Ecol. Lett. 24:1681–1696.

Heinhorst S., Cannon G. 1993. DNA Replication in Chloroplasts. J. Cell Sci. 104:1–9.

Hutchinson M.C., Cagua E.F., Balbuena J.A., Stouffer D.B., Poisot T. 2017. paco: implementing Procrustean Approach to Cophylogeny in R. Methods Ecol. Evol. 8:932–940.

Knowles L.L., Huang H., Sukumaran J., Smith S.A. 2018. A matter of phylogenetic scale: Distinguishing incomplete lineage sorting from lateral gene transfer as the cause of gene tree discord in recent versus deep diversification histories. American Journal of Botany. 105:376–384.

Krasnov B.R., Shenbrot G.I., Khokhlova I.S., Degen A.A. 2016. Trait-based and phylogenetic associations between parasites and their hosts: a case study with small mammals and fleas in the Palearctic. Oikos. 125:29–38.

Lagrue C., Joannes A., Poulin R., Blasco-Costa I. 2016. Genetic structure and host–parasite co-divergence: evidence for trait-specific local adaptation. Biol. J. Linn. Soc. 118:344–358.

Legendre P., Desdevises Y., Bazin E. 2002. A Statistical Test for Host–Parasite Coevolution. Syst. Bio. 51:217–234.

Léveillé-Bourret É., Eggertson Q., Hambleton S., Starr J.R. 2021. Cryptic diversity and significant cophylogenetic signal detected by DNA barcoding the rust fungi (Pucciniaceae) of Cyperaceae–Juncaceae. J Syst Evol. 59:833–851.

Pérez-Escobar O.A., Balbuena J.A., Gottschling M. 2016. Rumbling Orchids: How To Assess Divergent Evolution Between Chloroplast Endosymbionts and the Nuclear Host. Syst. Bio. 65:51–65.

Pérez-Escobar O.A., Dodsworth S., Bogarín D., Bellot S., Balbuena J.A., Schley R.J., Kikuchi I.A., Morris S.K., Epitawalage N., Cowan R., Maurin O., Zuntini A., Arias T., Serna-Sánchez A., Gravendeel B., Torres Jimenez M.F., Nargar K., Chomicki G., Chase M.W., Leitch I.J., Forest F., Baker W.J. 2021. Hundreds of nuclear and plastid loci yield novel insights into orchid relationships. Am. J. Bot. 108:1166–1180.

Pérez-Escobar O.A., Gottschling M., Chomicki G., Condamine F.L., Klitgård B.B., Pansarin E., Gerlach G. 2017. Andean Mountain Building Did not Preclude Dispersal of Lowland Epiphytic Orchids in the Neotropics. Sci Rep. 7:4919.

Poisot T. 2015. When is co-phylogeny evidence ofcoevolution? In: Krasnov B.R., Littlewood D.T.J., Morand S., editors. Parasite Diversity and Diversification: Evolutionary Ecology Meets Phylogenetics. Cambridge: Cambridge University Press. p. 420–433.

Raffinetti E., Siletti E., Vernizzi A. 2015. On the Gini coefficient normalization when attributes with negative values are considered. Stat Methods Appl. 24:507–521.

Renner S.S., Wu S., Pérez-Escobar O.A., Silber M.V., Fei Z., Chomicki G. 2021. A chromosome-level genome of a Kordofan melon illuminates the origin of domesticated watermelons. PNAS. 118:e2101486118.

Revell L.J. 2012. phytools: an R package for phylogenetic comparative biology (and other things). Methods Ecol. Evol. 3:217–223.

Rose J.P., Toledo C.A.P., Lemmon E.M., Lemmon A.R., Sytsma K.J. 2021. Out of Sight, Out of Mind: Widespread Nuclear and Plastid-Nuclear Discordance in the Flowering Plant Genus Polemonium (Polemoniaceae) Suggests Widespread Historical Gene Flow Despite Limited Nuclear Signal. Syst. Bio. 70:162–180.

Schardl C.L., Craven K.D., Speakman S., Stromberg A., Lindstrom A., Yoshida R. 2008. A Novel Test for Host-Symbiont Codivergence Indicates Ancient Origin of Fungal Endophytes in Grasses. Syst. Bio. 57:483–498.

Smith D.R., Arrigo K.R., Alderkamp A.-C., Allen A.E. 2014. Massive difference in synonymous substitution rates among mitochondrial, plastid, and nuclear genes of Phaeocystis algae. Mol Phylogenet Evol. 71:36–40.

Vargas O.M., Ortiz E.M., Simpson B.B. 2017. Conflicting phylogenomic signals reveal a pattern of reticulate evolution in a recent high-Andean diversification (Asteraceae: Astereae: Diplostephium). New Phytologist. 214:1736–1750.

