## Supplementary Material for "Rtapas: An R package to assess cophylogenetic signal between two evolutionary histories"

#### **Trematode and its amphipod host (Overview example)**

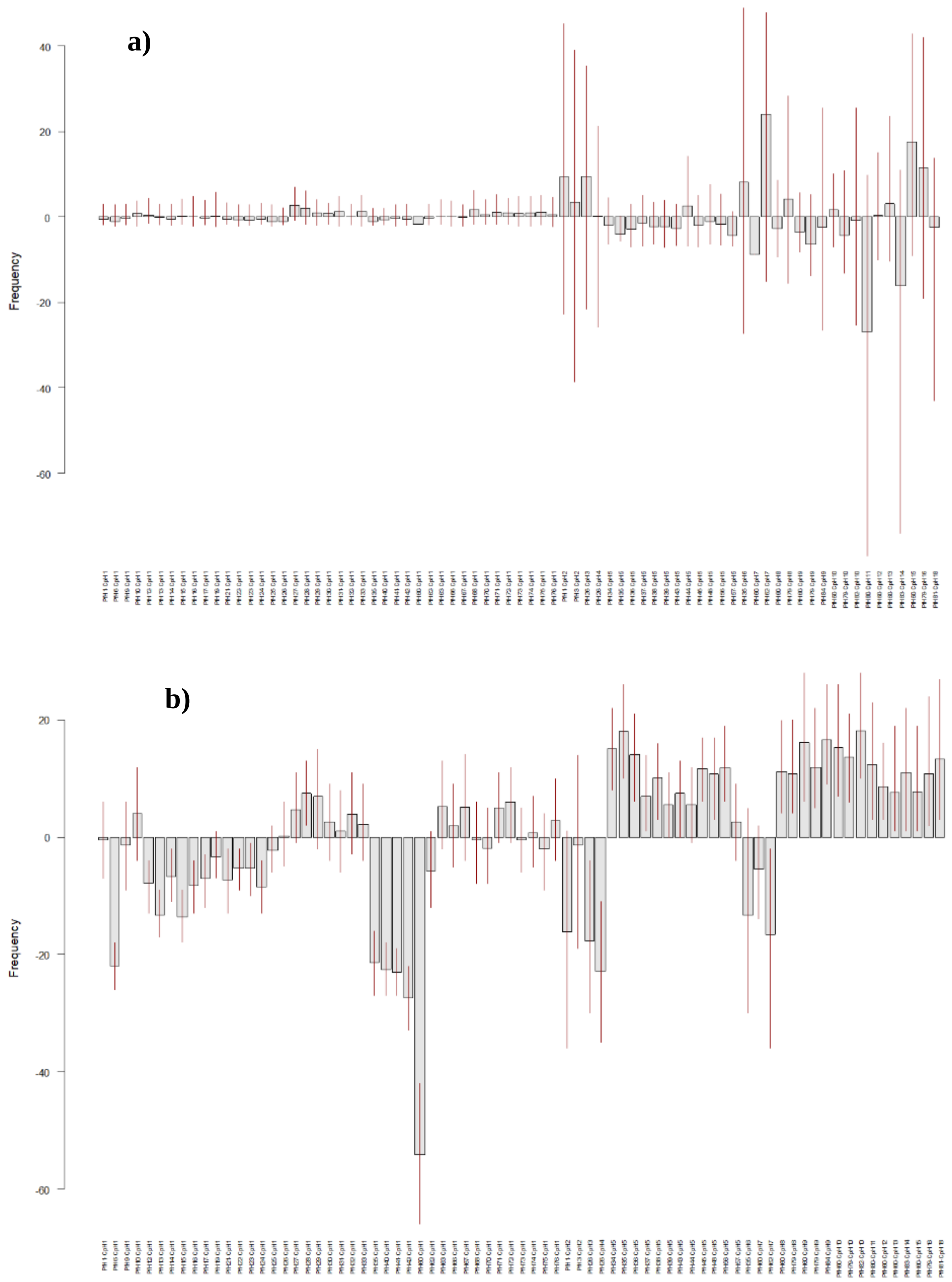


Figure S1. Average of (a) residual frequency distributions (observed – expected frequencies) for maximizing congruence algorithm, and (b) corrected frequency distributions for maximizing incongruence algorithm of host-symbiont associations of trematode and amphipod. Both algorithms were applied with PACo. Vertical lines represent 95% confidence intervals of the residual and corrected frequencies computed empirically with 1,000 randomly chosen pairs of posterior probability trees.

### Nuclear and plastid DNA data from orchids (Case Study 1)

This corresponds to the dataset of first case study in the manuscript. Here we provide the tanglegrams of the phylogenies corresponding to nuclear and plastid loci of orchids, obtained with the results of the frequencies presented in Figure 5 of the manuscript.

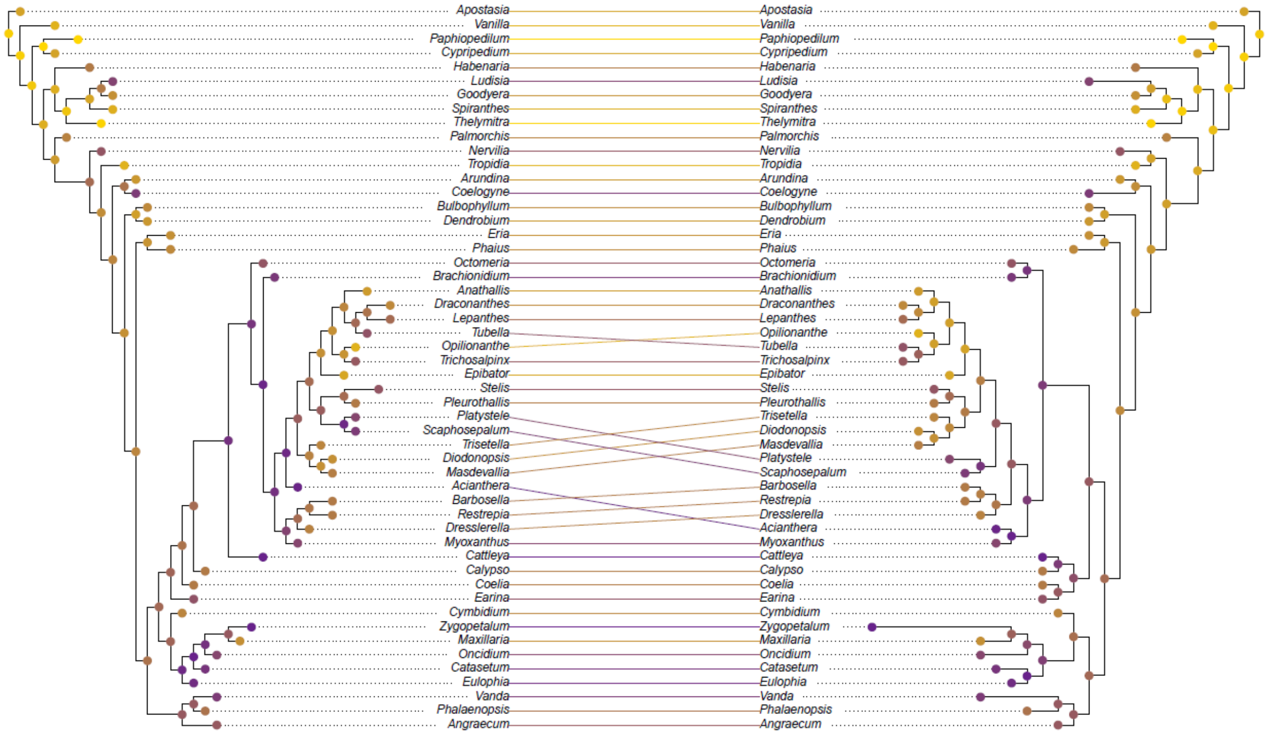


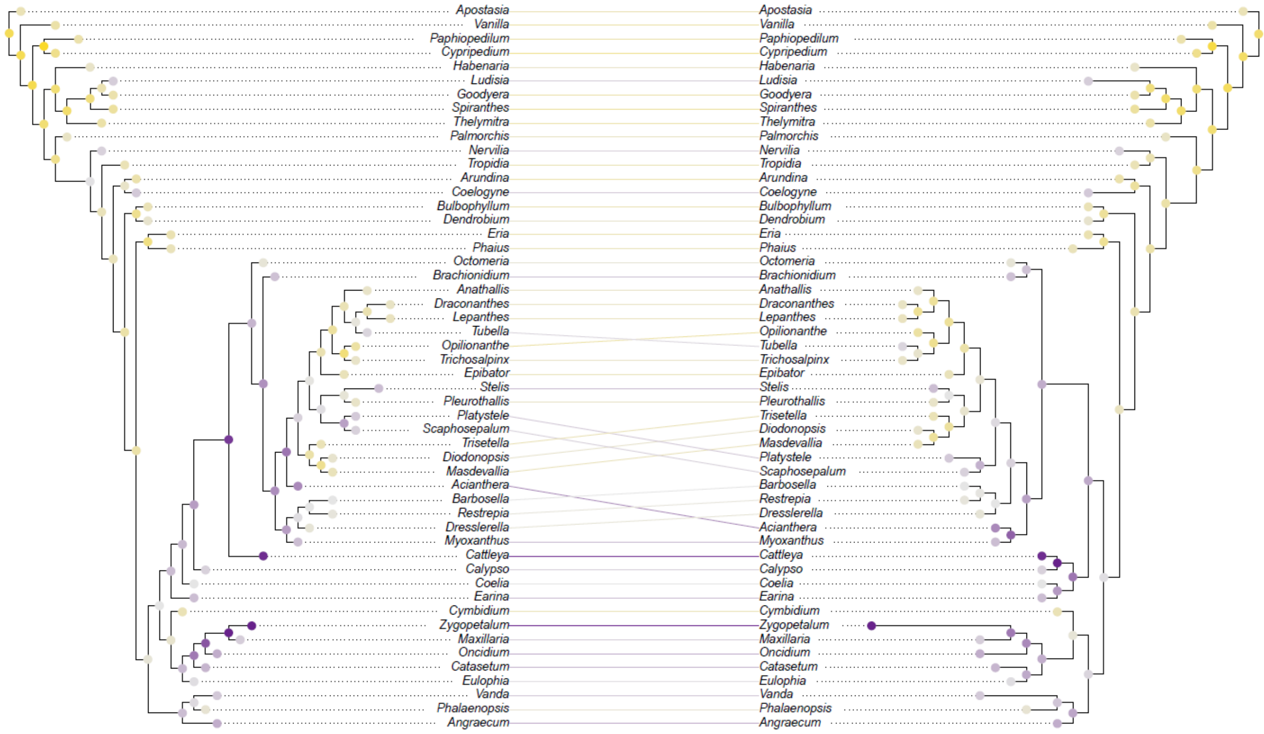

Figure S2. Tanglegrams representing the associations between organellar and nuclear DNA data from orchids. a) Displays the frequency corresponding to each plastid-nuclear association obtained with the algorithm to maximize congruence and b) displays the corrected frequency corresponding to each each plastid-nuclear association obtained with the algorithm to maximize incongruence. Both algorithms were applied with PACo. In a) the frequencies observed are mapped using a color ranging from the violet (minimum/incongruent) to gold (maximum/congruent). In b) the frequencies are centered at gray (zero) ranging from dark violet (lowest/incongruent) to gold (highest/congruent). The average frequency of occurrence of each terminal and fast maximum likelihood estimators of ancestral states of each node are also mapped according to the same scale, respectively.

#### Small mammals and their flea parasites (Case Study 2)

`Rtapas` can be applied to unravel complex patterns in large phylogenies. Can be used to observe generalized differences between several clades and, then, deepen the analysis within them. Here we break down the large phylogeny displayed in Figure 6 of the manuscript into three main clades (Eulipothyphla, Lagomorpha and Rodentia). This is achieved by trimming the association matrix with those species belonging to a given clade. Then we adjust the trees to match with the matrix (as seen in the following lines of code). Note that this is the procedure we follow, but other options are available. Once the data is trimmed, we apply the algorithm obtaining the residual and corrected frequencies, and displaying them as a heatmap in the tanglegrams. This dataset does not include posterior probabilistic trees, so we apply the algorithm without calculating the phylogenetic uncertainty. Even so, we believe that this analysis provides relevant information to interpret the ecological and speciation patterns of this system.

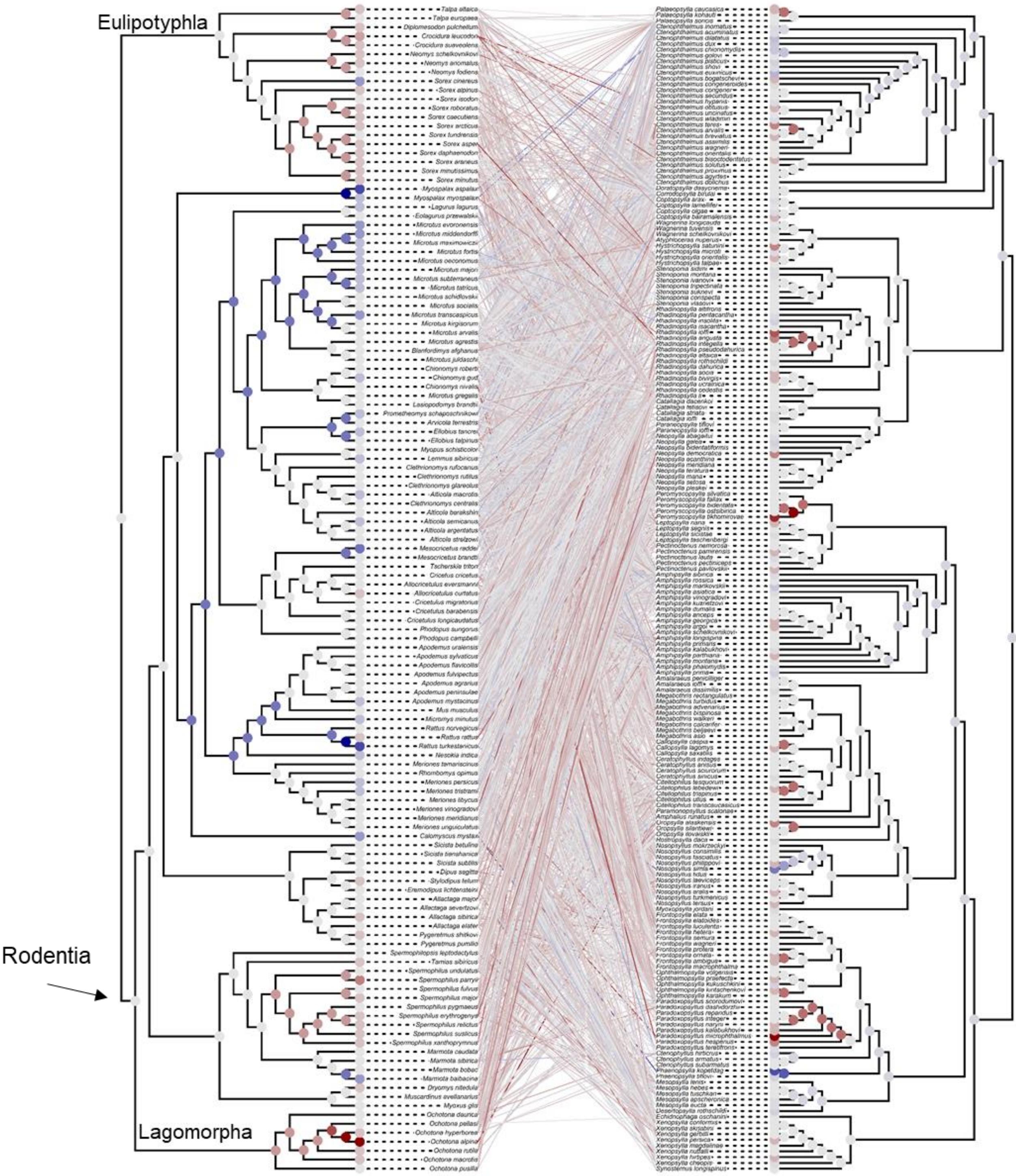


Figure S3. Tanglegram representing the associations between the mammal species with their flea parasite species with the algorithm to maximize congruence using PACo. The residual frequency (observed – expected) corresponding to each mammal-flea association obtained are mapped using a colour scale centred at light grey (zero) ranging from dark red (lowest/incongruent) to dark blue (highest/congruent). The average residual frequency of occurrence of each terminal and fast maximum likelihood estimators of ancestral states of each node are also mapped according to the same scale.

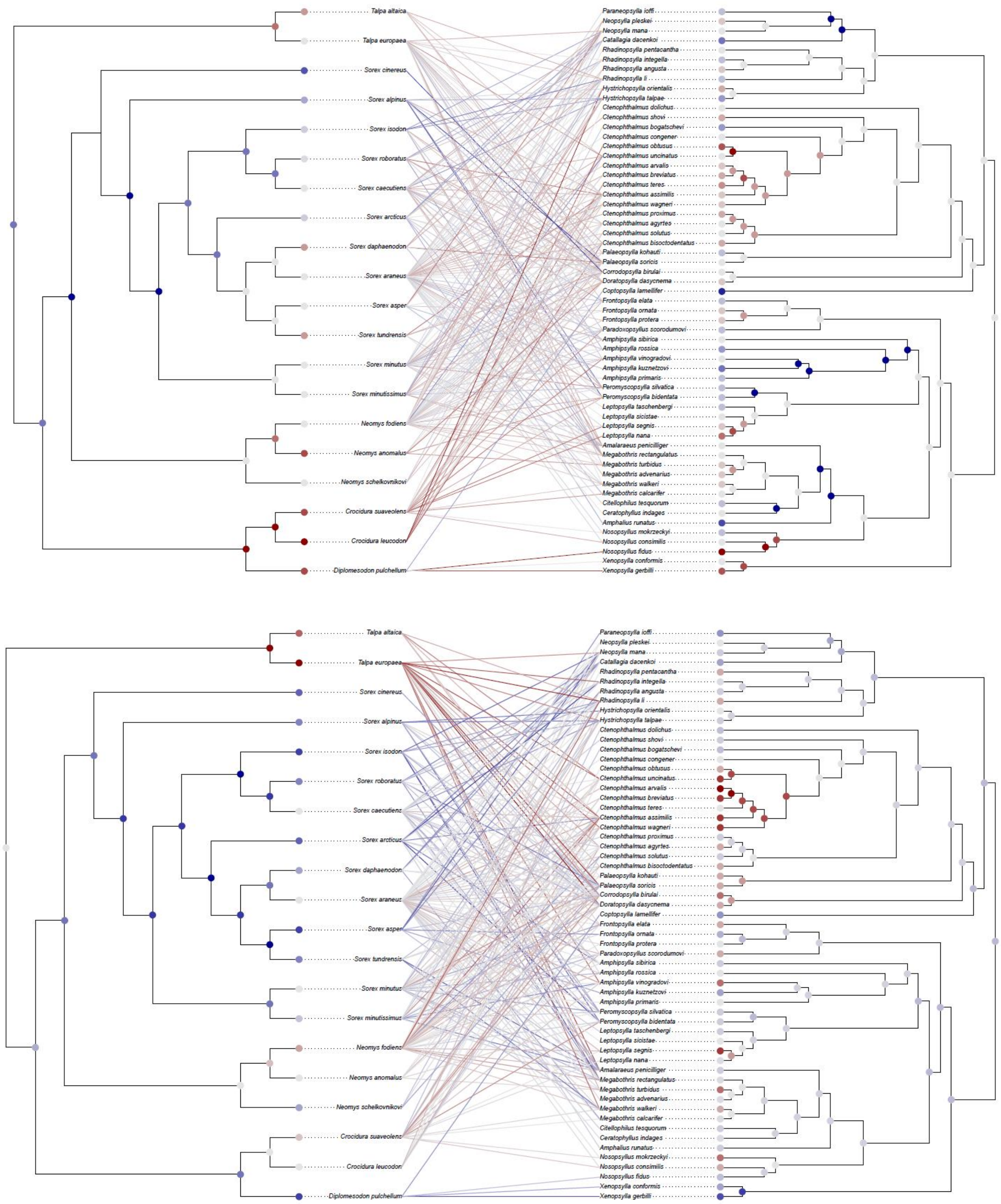


Figure S4. Tanglegrams representing the associations between mammals of the Eulipotyphla clade with their flea parasite species. a) Displays the residual frequency corresponding to each mammal-flea association obtained with the algorithm to maximize congruence and b) displays the corrected frequency corresponding to each mammal-flea association obtained with the algorithm to maximize incongruence. Both algorithms were applied with PACo. The frequencies are mapped using a color scale centered at light gray (zero) ranging from dark red (lowest/incongruent) to dark blue (highest/congruent). The average residual frequency of occurrence of each terminal and fast maximum likelihood estimators of ancestral states of each node are also mapped according to the same scale.

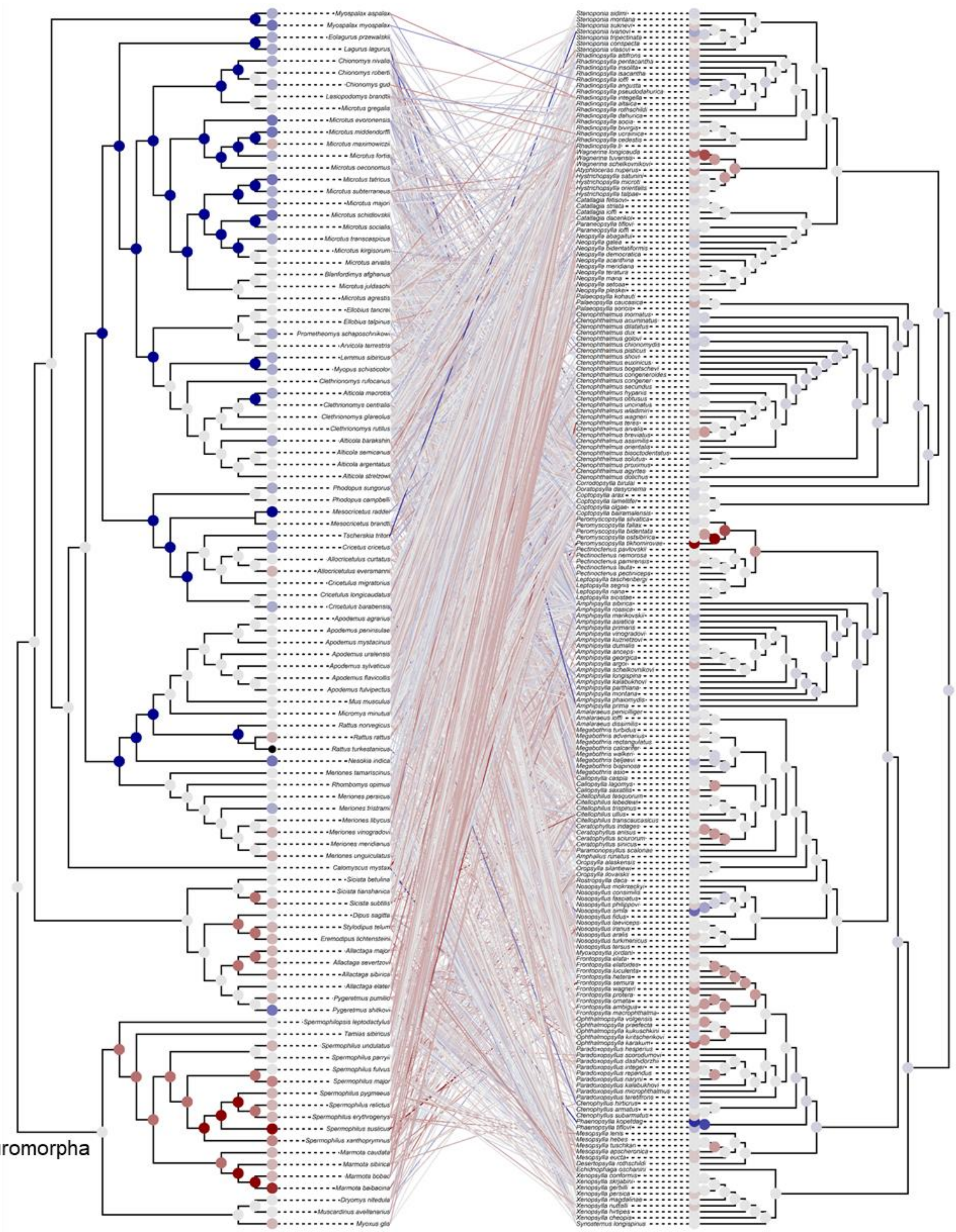


Figure S5a. Tanglegram representing the associations between mammals of the Rodentia clade with their flea parasite species with the algorithm to maximize congruence using PACo. The residual frequency (observed – expected) corresponding to each mammal-flea association obtained are mapped using a color scale centered at light gray (zero) ranging from dark red (lowest/incongruent) to dark blue (highest/congruent). The average residual frequency of occurrence of each terminal and fast maximum likelihood estimators of ancestral states of each node are also mapped according to the same scale.

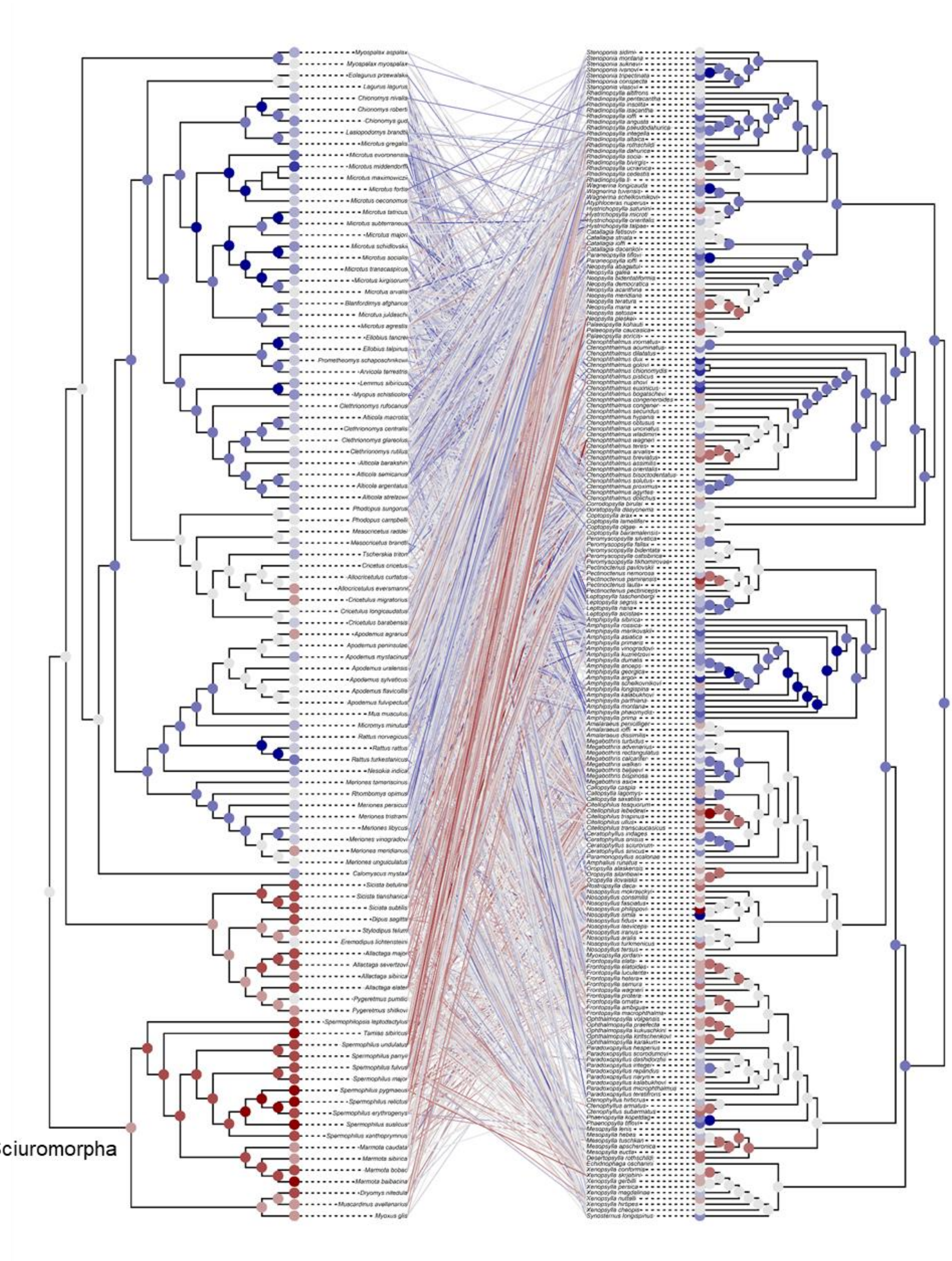


Figure S5b. Tanglegram representing the associations between mammals of the Rodentia clade with their flea parasite species with the algorithm to maximize incongruence using PACo. The corrected frequencies corresponding to each mammal-flea association obtained are mapped using a color scale centered at light gray (zero) ranging from dark red (lowest/incongruent) to dark blue (highest/congruent). The average residual frequency of occurrence of each terminal and fast maximum likelihood estimators of ancestral states of each node are also mapped according to the same scale.

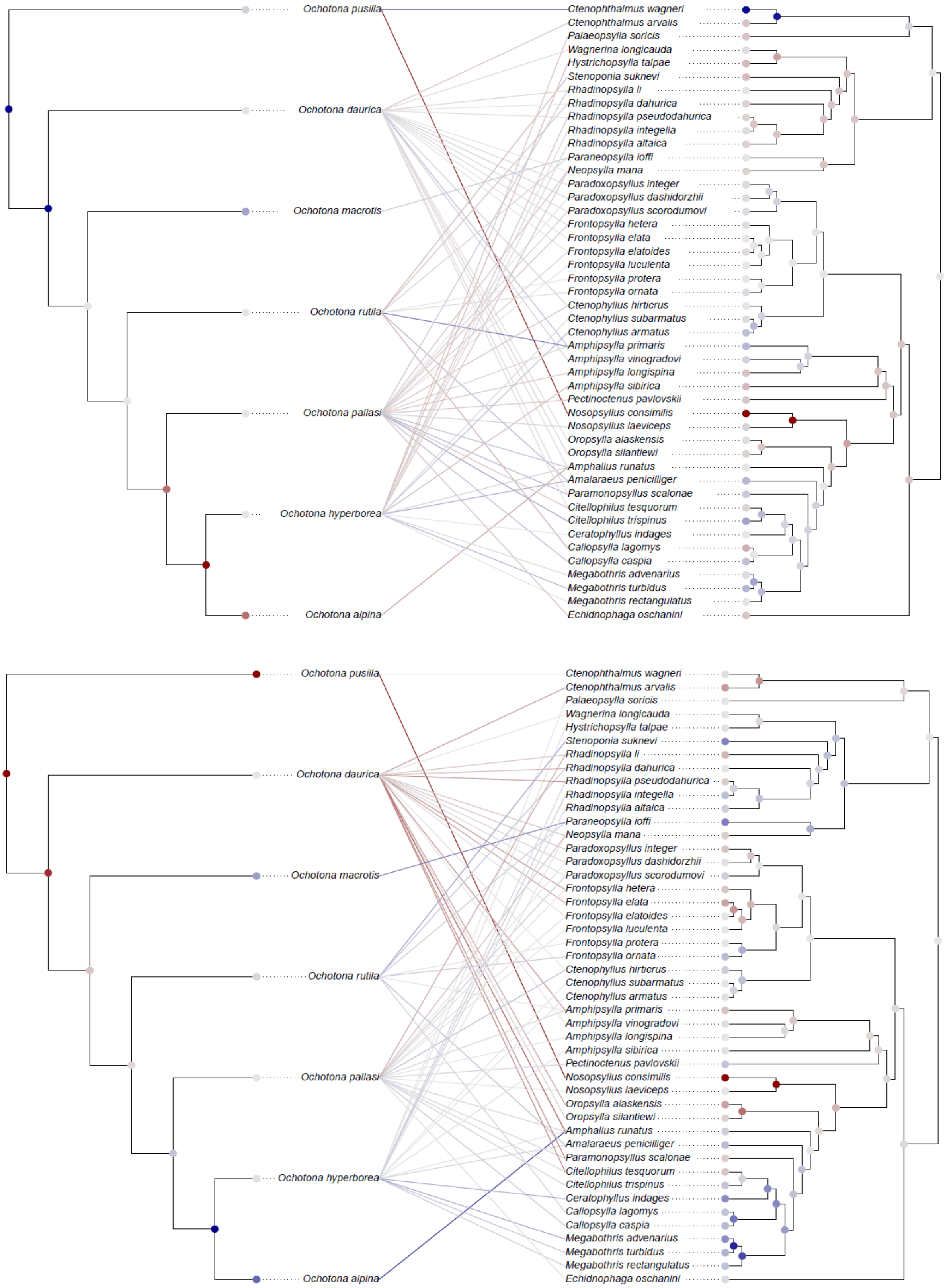


Figure S6. Tanglegrams representing the associations between mammals of the Lagomorpha clade with their flea parasite species. a) Displays the residual frequency corresponding to each trematode-amphipod association obtained with the algorithm to maximize congruence and b) displays the corrected frequency corresponding to each trematode-amphipod association obtained with the algorithm to maximize incongruence. Both algorithms were applied with PACo. The frequencies are mapped using a color scale centered at light gray (zero) ranging from dark red (lowest/incongruent) to dark blue (highest/congruent). The average residual frequency of occurrence of each terminal and fast maximum likelihood estimators of ancestral states of each node are also mapped according to the same scale.
